# The Conserved N-Terminal Extension of AtKEA1 Is Largely Dispensable for Plastid Function but Contributes to Potassium Homeostasis

**DOI:** 10.64898/2026.07.03.736308

**Authors:** Tobias Wunder, Lorenz Josef Holzner, Nikolay Manavski, Merve Nida Bastürk, Robert Janowski, Cäcilia Felicitas Kunz, Jonas Fechter, Susanne Mühlbauer, Florian Rösch, Jörg Meurer, Julia Legen, Franz Hagn, Dierk Niessing, Jan de Vries, Bettina Bölter, Hans-Henning Kunz

## Abstract

Members of the K⁺ efflux antiporter (KEA) family fulfill key roles in plastids and the endomembrane system. Plants and green algae possess at least one KEA mediating K⁺/H⁺ exchange across the plastid inner envelope (IE) membrane. Recently, IE KEAs were shown to be essential for plastid gene expression (PGE), chloroplast development, and photosynthesis. Plants lacking these antiporters exhibit reduced stromal protein synthesis and accumulation of unprocessed rRNA precursors. KEA proteins comprise a conserved monovalent cation/proton antiporter 2 (CPA2) domain and a regulatory K⁺ transport and NAD-binding (KTN) domain. IE KEAs are distinguished by an additional ∼500-amino-acid N-terminal extension containing a coiled-coil (CC) domain embedded within a largely intrinsically disordered region (IDR). Intrigued by this unusual architecture, we performed phylogenetic analyses, revealing that this N-terminal fusion arose early and has been conserved throughout the green lineage. We then investigated the oligomeric state, native distribution, and function of the N-terminal domain. Using *Arabidopsis thaliana*, we found that IE KEAs localize to discrete clusters within the inner envelope membrane and assemble into complexes of approximately 600 kDa. Finally, complementary approaches using a functional KEA1 variant lacking the core N-terminal domains (KEA1ΔN) indicate that this extension plays a regulatory rather than an essential role. Our findings uncover an evolutionarily ancient regulatory module that shapes the molecular organization and function of IE KEAs, advancing our understanding of plastid ion and pH homeostasis and plastid ribosome integrity.

**One-sentence summary:** Plastid KEA1/2 proteins feature a unique N-terminal extension that modulates potassium transport activity in a yet unknown manner but is not essential for normal plant growth under ambient conditions.

## Introduction

Chloroplasts serve as the central hub for photosynthesis and numerous biosynthetic pathways essential for plant growth and development. The proper functioning of these processes depends on the establishment and maintenance of ion homeostasis across chloroplast compartments, which underpins both bioenergetic and metabolic activities (Kunz et al., 2024). Within this context, the K⁺/H⁺ exchanger (KEA) proteins of *Arabidopsis thaliana* have emerged as key regulators of plastid potassium balance and pH control. Belonging to the cation/proton antiporter 2 (CPA2) superfamily, the KEA family comprises six members with distinct localizations and functions. KEA1 and KEA2 are localized to the inner envelope (IE) membrane of the chloroplast and form the KEA 1a clade (Kunz et al., 2014; Chanroj et al., 2012), whereas KEA3, a clade 1b protein (Chanroj et al., 2012), resides in the thylakoid membrane, where it fine-tunes lumenal pH during photosynthetic light responses (Armbruster et al., 2014; Kunz et al., 2014).

A distinctive and phylogenetically unique feature of the IE-localized KEAs is their extended N-terminal domain, which protrudes into the plastid stroma (Bölter et al., 2020). This soluble region is absent from other CPA2 family members and contains characteristic structural elements. However, both its evolutionary origin and functional role remain unclear. Elucidating the evolutionary history of this domain in the context of the transporter module is therefore essential for understanding its biological significance.

Functional studies have shown that simultaneous loss of KEA1 and KEA2 in *kea1kea2* double loss-of-function mutant plants leads to severe developmental defects, particularly in young, expanding leaves (Kunz et al., 2014). These defects include impaired chloroplast biogenesis, disrupted thylakoid architecture, reduced chlorophyll accumulation, and compromised plastid gene expression (PGE) (DeTar et al., 2021; DeTar et al., 2022). The resulting ionic imbalance is thought to disturb the electrostatic conditions required for proper ribosome processing, assembly, and function within the stroma.

Another largely unexplored aspect of IE KEA biology is their oligomeric organization within the inner envelope membrane. Many plastid membrane proteins form high-molecular-weight complexes (Lundquist et al., 2017), yet the native assembly state of KEA1 and KEA2, as well as the molecular determinants governing their complex formation, remain unknown.

In this study, we address these questions using a combination of phylogenetic, biochemical, cell biological, and molecular approaches. We first investigated the evolutionary origin and structural features of the IE KEA N-terminal domain. We then established complementation lines expressing fluorescently tagged full-length KEA1 and an N-terminally truncated variant (*KEA1ΔN*) under the native promoter in the *kea1kea2* background, enabling a systematic analysis of the N-terminus in protein localization, oligomerization, potassium transport, and photosynthetic performance. Finally, we examined rRNA processing in *kea1kea2* mutants and the complemented lines to assess downstream effects on plastid function.

## Results and Discussion

### Evolution of the Inner Envelope KEAs is linked to a unique, soluble N-terminus absent from other related ion transporters

IE KEA transporters, KEA1 and KEA2, are essential for plastid ion homeostasis (Fig. 1A). Like other members of the CPA2 family, they possess a conserved core architecture consisting of a transmembrane K⁺/H⁺ antiporter domain and a regulatory C-terminal nucleotide-binding (KTN) domain. What distinguishes KEA1 and KEA2 from all other CPA2 family members is their extended N-terminal domain, which reaches into the plastid stroma (Fig. 1A) (Bölter et al., 2020). This additional soluble segment of more than 500 amino acids contains a coiled-coil (CC) domain (Zhang and Dang, 2026) embedded within a largely intrinsically disordered region (IDR) rich in basic and acidic residues (Fig. 1B).

**Figure 1.**
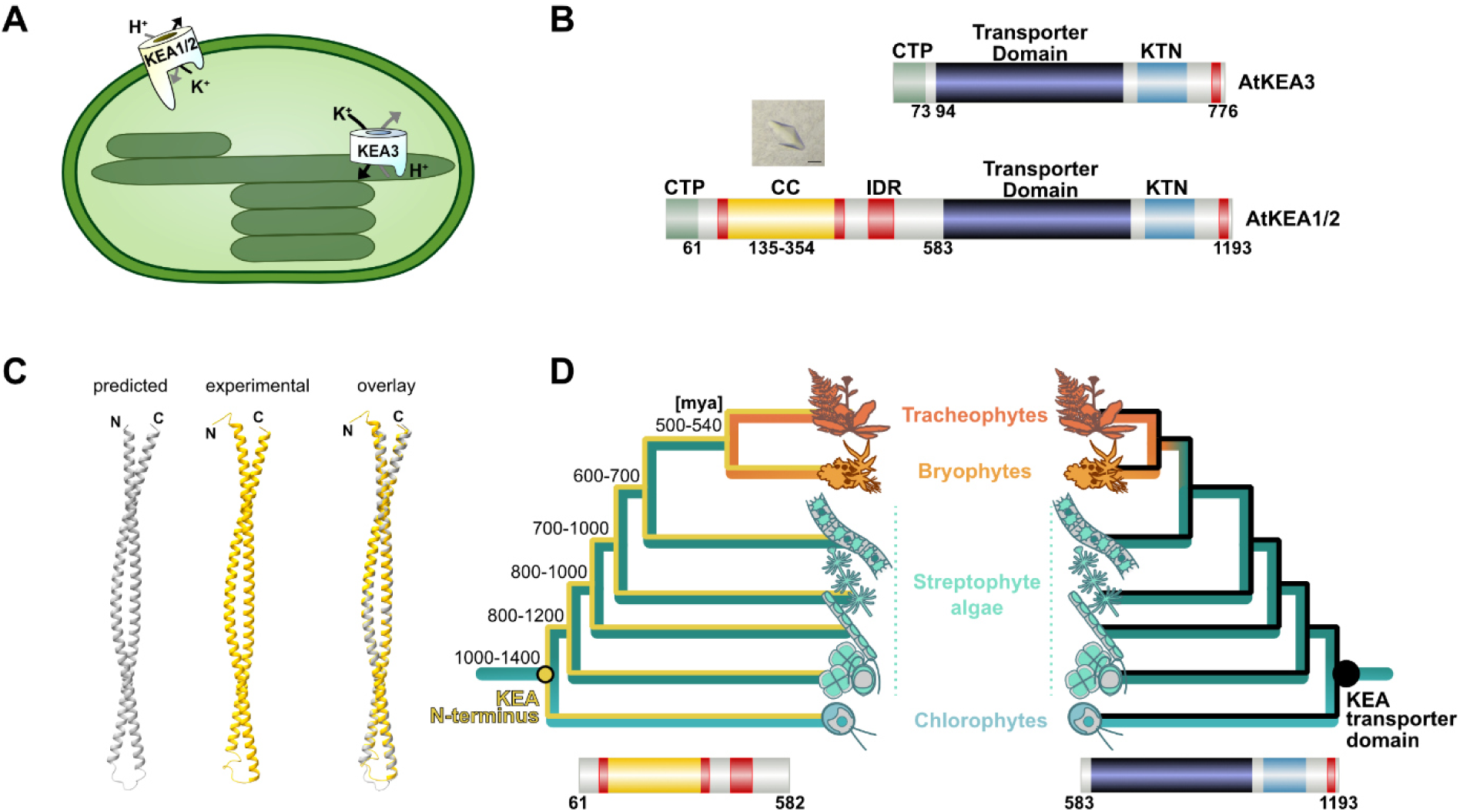
KEAs of the plastid inner envelope (IE) membrane feature a unique N-terminus. **A)** Simplified model of plastid KEAs’ location and function. **B)** Schematic domain organization of the inner-envelope transporters KEA1/2 and the thylakoid-localized KEA3 based on UniProt annotations (Consortium, 2025): coiled-coil domain (CC) of KEA1/2; CTP, chloroplast targeting peptide; IDR, intrinsic disordered regions; KTN, Potassium Transport–Nucleotide binding domain. Crystal of the KEA1 CC (aa 135–354) is shown above the respective domain. Scale bar = 0.1 mm. **C)** Predicted (grey) and experimental (yellow) structure of the CC fragment determined by X-ray crystallography at 3.59 Å resolution, shown in overlay (gray/yellow). **D**) Cladogram of Chloroplastida, including chlorophytes (blue), streptophyte algae (turquoise; from bottom to top: mesostigmatophytes and chlorokybophytes, klebsormidiophytes, charophytes, and zygnematophytes), and embryophytes (orange, bryophytes; red, tracheophytes). The left panel depicts the phylogeny of KEA1’s N-terminus (yellow), whereas the right panel shows the evolution of the KEA1 transporter domain (black).

To gain structural insight into the function of the N-terminal extension, we pursued purification (Fig. S1A) and X-ray crystallographic analysis. Because large portions of the N-terminus are predicted to be intrinsically disordered, we aimed at solving the crystal structure of the putative CC region. From the KEA1(135–354) fragment, we obtained a single crystal (Fig. 1B, inset above the CC domain) that diffracted to 3.59 Å resolution. The asymmetric unit contains two copies of KEA1(135–354) (Fig. S1B), and the crystal exhibits an unusually high solvent content (86%; Matthews coefficient 8.6), which likely contributed to the limited diffraction quality. Although optimization did not improve crystal quality, the dataset with 3.59 Å resolution was sufficient to build a preliminary model. While side-chain assignment was not possible, the electron density clearly defined the protein backbone (Fig. S1C) and confirmed the predicted coiled-coil architecture (Fig. 1C). Analysis of crystal contacts revealed two possible dimeric arrangements, one parallel and one antiparallel (Fig. S1D). However, given the crystal packing environment, we cannot exclude the possibility that these interactions represent crystallographic rather than physiological interfaces. Overall, the structure provides experimental proof for the predicted CC motif in the KEA1 N-terminal region. The occurrence of IE KEA featuring the N-terminal extension is restricted to eukaryotic photosynthetic organisms that fall into the green lineage, the Chloroplastida. The N-terminal extension is equally present in representatives of the Chlorophyta and the Streptophyta, the latter including land plants and the paraphyletic streptophytic green algae (Kunz et al., 2025; Basturk et al., 2026). IE KEAs were not detected in the other two Archaeplastidial lineages, Rhodophyta and Glaucophyta, or in Prasinodermatophyta, the proposed sister lineage to Chlorophyta and Streptophyta (Li et al., 2020). Although *Prasinoderma coloniale* contains K4/5/6 and KEA3 homologs, KEA1/2 homologs are absent.

Therefore, we hypothesize that the fusion of KEA to its soluble N-terminal extension evolved to serve a functional role in the green lineage, making it an important synapomorphy of Chloroplastida. We next examined whether the distinctive N-terminus of IE KEA proteins was acquired through repeated independent events during evolution or originated from a single ancestral fusion.

To address this, we generated phylogenetic trees and co-phylogeny plots for pairwise comparisons of IE KEA domains (Fig. 1D, Fig. S1E). When comparing the phylogenetic tree of the N-terminus (KEA1 residues 61–582) with that of the transporter domain lacking the N-terminus (KEA1 residues 583–1193), the overall topologies showed minimal differences. Despite the faster divergence in the primary sequence of the N-terminus, its evolutionary trajectory closely mirrors that of the transporter domain. This strongly suggests that a single, early fusion event occurred, linking the transporter domain (physically and evolutionarily) to its N-terminal extension. A lack of structural constraints usually facilitates faster evolution at the amino acid sequence level (Brown et al., 2011). Consequently, the largely disordered N-terminus evolved more rapidly than the structurally conserved transporter and KTN domains.

In summary, IE KEAs possess a unique and distinctive long N-terminal extension. This domain, which includes a CC motif within an IDR, appears to have been acquired early during the evolution of the green lineage (≳1 billion years ago) through a single evolutionary event. Despite rapid divergence at the amino acid level, the structural conservation of this N-terminal region suggests a critical functional role. While the precise function remains elusive, its involvement in regulatory or organizational processes within the chloroplast stroma is strongly implied by the structural stability and the well-documented roles of coiled-coil motifs in facilitating protein-protein interactions (Kodama et al., 2011; Seung et al., 2018).

### KEA1ΔN functionally complements *kea1kea2* and accumulates in discrete spots in the plastid inner envelope

Before investigating the function of KEA1’s intriguing N-terminus in more detail, we established an expression system that allowed us to study the protein under close to native conditions.

In a previous study, KEA1/2 and the single homolog from *Oryza sativa* OsKEA1 were either observed in specific spots in the plastid envelope membrane or distributed uniformly throughout the envelope (Kunz et al., 2014; Aranda-Sicilia et al., 2016). Additionally, KEA2 accumulated in the pole region of developing chloroplasts (Aranda-Sicilia et al., 2016). These studies employed the constitutive promoters *AtUBQ10* (*pUBQ10*) or *CaMV 35S*, respectively, to drive IE KEA expression. However, their high and ubiquitous expression levels may conceal more nuanced spatial expression patterns (Kunz et al., 2014; Aranda-Sicilia et al., 2016). Therefore, we exchanged *pUBQ10* for the endogenous *KEA1* promotor (*pKEA1*) and focused our research on KEA1, previously identified as the predominant isoform expressed in Arabidopsis leaves (Bölter et al., 2020). To test their functionality, the constructs expressing YFP-tagged *KEA1* and *KEA1^480-1193^*Δ*N* (aa 1-100 containing the chloroplast targeting information fused to aa 480-1193; henceforth labelled as *KEA1-YFP* and *KEA1*Δ*N-YFP,* respectively) were inserted into the genome of the double mutant *kea1kea2* (Kunz et al., 2014). As previously observed for the *pUBQ10*-driven lines (Fig. S2A,B) (Bölter et al., 2020), *KEA1-YFP* transformants were comparable to wild type (WT) control plants with regards to their appearance and the maximum quantum efficiency of photosystem II (PSII) photochemistry (*F*_v_/*F*_m_) (Fig. 2A). Intriguingly, the same was true for the *KEA1*Δ*N-YFP* line, indicating that the highly conserved N-terminus is not necessary for general KEA1 function under the employed ambient growth conditions. Transgene expression was validated by immunoblot analysis using an α-GFP antibody alongside the previously characterized α-KEA1 antiserum (Bölter et al., 2020), confirming that protein accumulation levels of KEA1-YFP and KEA1ΔN-YFP were comparable and similar (slightly higher) to those observed in WT plants (Fig. 2B, Fig. S2B). Since the KEA1 antibody was raised against the N-terminus, KEA1ΔN-YFP is detectable exclusively with the GFP antibody.

**Figure 2.**
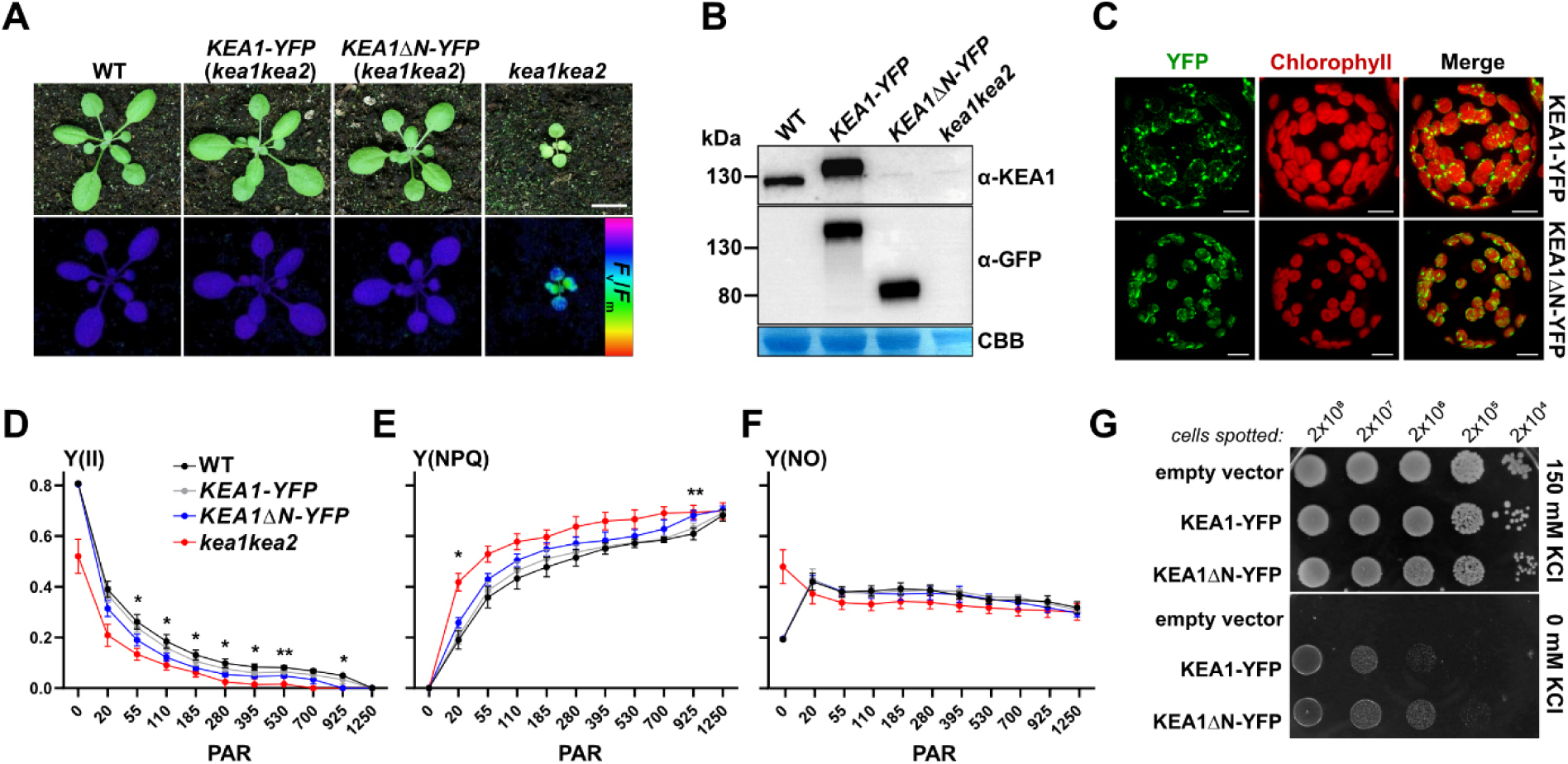
The N-terminal extension of KEA1 is not essential for potassium transport activity. **A)** RGB and false-color *F*_v_/*F*_m_ images of WT, *kea1kea2* null mutants, and complementation lines expressing KEA1-YFP or KEA1ΔN-YFP under the control of the native KEA1 promoter in the *kea1kea2* background. Scale bar = 1 cm. **B)** Immunoblot using AtKEA1 and GFP antibodies to detect KEA1 expression in the complementation lines. Coomassie-stained (CBB) RbcL (50–60 kDa region) as a loading control. **C)** Confocal laser scanning microscopy of protoplasts reveals a punctate distribution of KEA1-YFP and KEA1ΔN-YFP surrounding the chloroplast chlorophyll autofluorescence. Scale bar = 10 µm. **D–F)** PAM measurements during a standard light curve. Y(II), effective quantum yield of PSII; Y(NPQ), non-photochemical quenching; Y(NO), non-regulated energy dissipation. Data are means ± SD (n = 6 biological replicates). Statistical analysis was performed by two-way repeated-measures ANOVA with Geisser-Greenhouse correction, followed by Tukey’s multiple-comparisons test. Asterisks indicate p values for the KEA1-YFP vs KEA1ΔN-YFP comparison only (*p<0.05, **p<0.01p, ***p<0.001, ****p<0.0001). **G)** Both KEA1-YFP and KEA1ΔN-YFP complement the potassium transport-deficient *E. coli* strain LB2003 under potassium-limiting conditions.

To study subcellular localization, we imaged protoplasts isolated from both complementation lines. Fluorescence microscopy revealed distinct YFP signals in both cell lines, with KEA1 and KEA1ΔN localizing to discrete punctate subdomains within the IE, consistent with previously reported findings for KEA1 (Fig. 2C) (Kunz et al., 2014; Aranda-Sicilia et al., 2016). While in our study cluster formation was not abolished in *KEA1ΔN* lines, a truncation of KEA2 - albeit ∼100 amino acids shorter than the construct used here - resulted in a more uniform distribution around chloroplasts (Aranda-Sicilia et al., 2016). When we generated a similarly truncated *KEA1^578-1193^ΔN–YFP* construct (aa 1-60 fused to aa 578-1193, Fig. S3A), no phenotypically complemented hygromycin-resistant plants were identified (Fig. S3B). These observations suggest that the region between aa 480-578, absent in the shorter KEA1^578-1193^ΔN-YFP version, is essential for proper targeting, membrane insertion, and/or function of IE KEA proteins.

As the punctate distribution pattern was reminiscent of nucleoid distribution (Powikrowska et al., 2014; Legen et al., 2024), we conducted STED microscopy on DAPI-stained protoplasts from WT plants and the *KEA1–YFP* line. When we carefully examined the plastid DAPI and YFP signals, we found no evidence for preferential spatial association between KEA1 and nucleoids (Fig. S4A). Specifically, distance distribution analysis did not reveal a distinct peak. Thus, the punctate localization of KEA1 cannot be readily attributed to nucleoid association, suggesting that the observed puncta reflect another aspect of KEA1 organization within the chloroplast envelope (Fig. S4B).

Photosynthetic parameters measured during light curves, Y(II), Y(NPQ), and Y(NO), the latter representing non-regulated energy dissipation in (PSII) (Kramer et al., 2004), closely resembled WT levels in KEA1. In contrast, the *KEA1ΔN-YFP* line showed a modest but consistent reduction in PSII photochemical efficiency (Y(II)), accompanied by slightly increased non-photochemical quenching (Y(NPQ)), while Y(NO) remained largely unchanged across the light range (Fig. 2D–F). These shifts indicate a subtle but reproducible redistribution of excitation energy toward regulated dissipation (NPQ), suggesting a minor functional impairment in photosynthetic energy partitioning rather than a pronounced physiological defect.

To further evaluate the general potassium transport activity of KEA1-YFP and KEA1ΔN-YFP, the corresponding cDNAs with bacteria-optimized codon-usage were subcloned into the pBAD expression vector and expressed in the potassium transport-deficient *E. coli* strain LB2003 (Stumpe and Bakker, 1997) which has successfully been applied to demonstrate potassium transport activity of KEA1-3 (Tsujii et al., 2019). Complementation was assessed by spotting serial dilutions (2 × 10⁸ to 2 × 10⁴ cells) onto LB plates not supplemented with potassium and comparing growth restoration to that on potassium-supplemented plates (Fig. 2G). Whereas the empty vector failed to rescue the growth phenotype in the absence of supplemented potassium, both constructs partially restored growth under these conditions. Notably, KEA1ΔN-YFP complemented the mutant more efficiently than full-length KEA1 (Fig. 2G), supporting a modulatory role of the N-terminal domain in transporter activity, at least in the heterologous *E. coli* system lacking chloroplast-specific regulatory factors (Fig. 2G). This interpretation is further supported by the subtle but reproducible alterations in photosynthetic energy partitioning observed in the *KEA1ΔN* complementation line, which likewise point to a regulatory function of the N-terminal domain under physiological conditions.

While heterologous complementation assays are consistent with the N-terminal domain exerting a negative effect on transport activity (Fig. 2G) (Aranda-Sicilia et al., 2012; Zheng et al., 2013), the physiological role of this domain in planta may be more complex. The observation that KEA1ΔN largely restores growth and plastid function raises the possibility that the N-terminus contributes to context-dependent regulation of KEA activity rather than acting solely as an inhibitory module. Thus, deletion of the N-terminus may uncouple transport activity from native regulatory inputs, yielding a functional transporter that fails to fully restore wild-type ion balance.

### IE KEAs assemble into large oligomers independent of the N-terminus

Most plastid proteins reside in high-molecular-weight complexes *in vivo* (Lundquist et al., 2017). To assess the native oligomeric state of KEA1 and KEA2 within the distinct IE clusters observed in confocal microscopy (Fig. 2C), we employed blue native (BN) PAGEs and size exclusion chromatography (SEC).

Initially, we isolated chloroplasts from WT, *kea1kea2*, and *kea3* (*kea3-1* allele, (Kunz et al., 2014)) lines. KEA3 (∼70 kDa monomeric) was recently described to oligomerize *in vivo* (Uflewski et al., 2021). Consistent with this, KEA3 was detected in a ∼150 kDa complex in our BN-PAGE (Fig. 3A), but not in higher order complexes as previously suggested by Uflewski.

**Figure 3.**
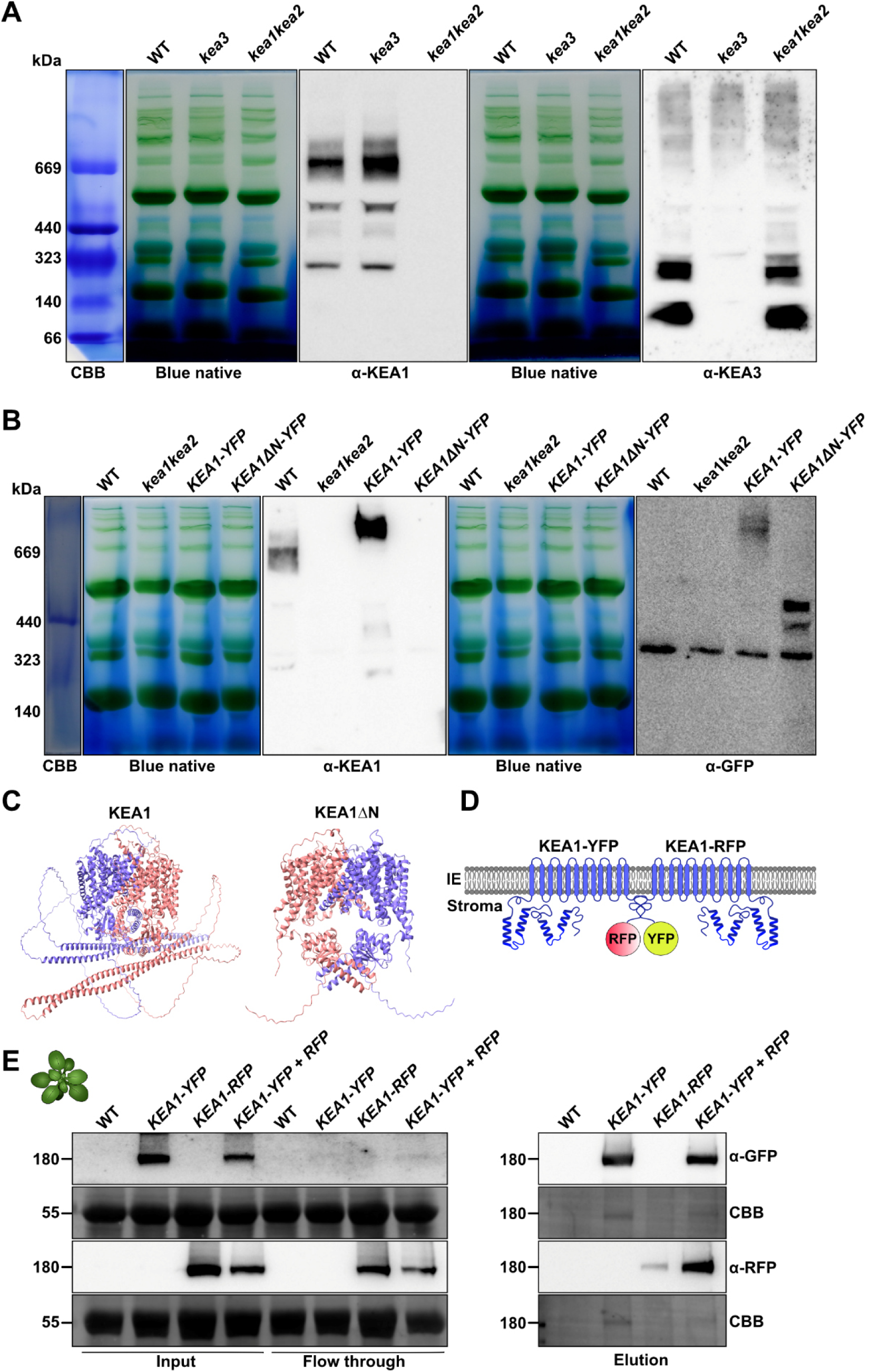
KEA1 self-associates into protein complexes. A, **B)** Blue Native-PAGE analysis of detergent-solubilized chloroplasts from the indicated genotypes. **C)** AlphaFold3-predicted structures of mature KEA1 variants modeled as homodimers. **D)** Schematic representation of two fluorescence-tagged KEA1 monomers interacting via their C-termini. **E)** GFP-Trap immunoprecipitation demonstrates co-purification of RFP-tagged KEA1 with KEA1-YFP from the double-expression line.

For KEA1 (∼123 kDa calculated monomeric size), most of the protein was found in a complex migrating around 660 kDa in the BN-PAGE. Additionally, weaker signals were detected above 660 kDa and at approximately 500 kDa and 230 kDa (Fig. 3A). Note that membrane protein complexes tend to show higher than expected apparent molecular weights (Ma and Xia, 2008). Thus, the true molecular weight cannot be reliably estimated from BN-PAGE mobility.

Therefore, chloroplast protein extracts were separated by SEC in parallel (Fig. S5A). Subsequently, the collected fractions were immunoblotted using α-KEA1 and α-KEA3, respectively. KEA1 peaked in the 13 – 14 ml fraction that corresponds well to a size of approximately 660 kDa. For KEA3, one fraction eluted well after the 159 kDa standard (Fig. S5A), approximately matching the 120 kDa result obtained from BN analysis, while a second peak can be detected in the 15 – 16 ml fractions, indicating the presence of a larger complex of ∼350 kDa that was not detectable in the BN-PAGE.

Since CC motifs are well known to mediate protein dimerization and higher-order assembly (Maekawa et al., 2011), we investigated whether this domain is required for IE KEA oligomerization. To this end, we employed the same BN-PAGE approach to assess the complex formation status of the *KEA1ΔN* truncation line, in which the N-terminal region with its coiled-coil domain is absent. Immunoblot analysis of BN-PAGE-separated chloroplast protein extracts revealed that the *KEA1ΔN* line retains the ability to from higher-order complexes with an apparent molecular weight of ∼400 kDa, while the monomer runs at 90 kDa in SDS-PAGE (Fig. S5B,C). These complexes migrated at a lower apparent molecular weight than those of full-length KEA1, consistent with the reduced size of the truncated protein (Fig. 3B, right panel).

Together, these findings suggest that while the N-terminal coiled-coil domain may contribute to the stability or stoichiometry of KEA1 complexes, it is not strictly required for oligomerization. This observation is further corroborated by AlphaFold3 (Abramson et al., 2024) structural modelling of both KEA1 and KEA1ΔN-YFP dimers (Fig. 3C), which predicts that dimerization occurs independently of the N-terminal CC domain. Instead, the dimeric interface appears to be primarily mediated by the C-terminal KTN domain, with a potential additional contribution from select channel-forming transmembrane α-helices. These structural predictions are consistent with the biochemical data and suggest that the KTN domain serves as the principal determinant of KEA1 dimerization, a mechanism that has been described for other KTN domain-containing transporters (Roosild et al., 2002).

To investigate the molecular composition of IE KEA-containing complexes, GFP-TRAP–based pulldown assays were performed (Schwartz et al., 2025) (Fig. 3D). Complemented Arabidopsis *kea1kea2* lines expressing *KEA1-YFP or KEA1-RFP* under the native promoter were crossed, and protein expression was verified by immunoblotting (Fig. S5D).

Our pulldown results show that KEA1-RFP can be precipitated with co-expressed KEA1-YFP (Fig. 3E), indicating that the previously observed complex of IE KEAs contains at least two KEA1 subunits. These results were confirmed by transiently expressing tagged KEA1 proteins in tobacco leaves (Fig: S5E-G). Utilizing the transient *N. benthamiana* system, we also found that RFP-tagged KEA2 was detectable in KEA1-YFP precipitates when both constructs were expressed simultaneously. It follows that the IE KEA complexes could theoretically consist of KEA1-KEA2 hetero-oligomers. While KEA2 was previously found to accumulate only to a low protein level (Bölter et al., 2020), the relative contributions of either isoform may vary depending on the physiological or developmental state of the tissue. Notably, heteromeric formation seems not essential, as single null mutants of either KEA1 or KEA2 display phenotypes indistinguishable from the WT (Kunz et al., 2014). Also, other eukaryotic photosynthetic organisms like the green alga *Chlamydomonas* (Wunder et al., 2026), but also the moss *Physcomitrium* and vascular plants like rice (Sheng et al., 2014), contain only a single locus encoding for the IE KEA homolog (Chanroj et al., 2012).

Taken together, pulldown and co-precipitation data collectively support the existence of plastid protein complexes containing IE KEA proteins. To date, no additional interaction partners have been identified, though their presence cannot be ruled out.

### *kea1kea2* mutants accumulate polysome-associated 23S-4.5S rRNA precursors

Characterization of the *kea1kea2* double mutants revealed impaired ribosome maturation (DeTar et al., 2021), prompting a comparative analysis of our transgenic lines. RNA blots using five different probes (A-E) binding along the 23S, 4.5S, and 5S rRNAs, respectively, were carried out. Only the *kea1kea2* genotype showed a clear accumulation of the largest detectable RNA species, the 3.2 kb 23S - 4.5S precursor, which is recognized by probes A-C targeting the 23S region and probe D targeting the 4.5S region (Figure 4A,B). In addition to the presence of the unprocessed 3.2 kb fragment, probe D also revealed a clear decrease in mature 4.5S i.e., the 100 nt fragment. In contrast, probe E specific for the 5S fragment showed no difference in abundance compared to WT extracts, indicating that the effects observed in *kea1kea2* plastids are upstream of the 5S processing and thus very specific towards the 23S – 4.5S rRNA bicistronic precursor (Figure 4B).

**Figure 4.**
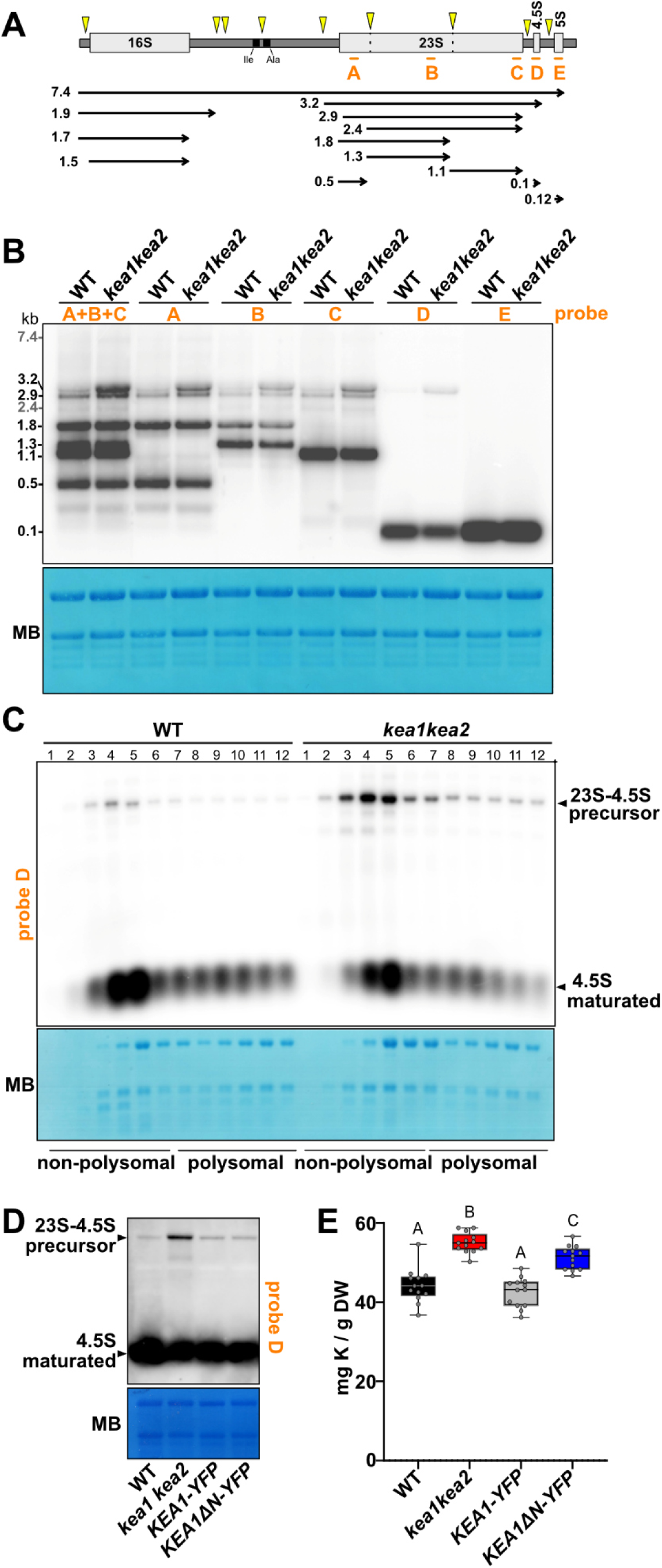
KEA1 lacking its N-terminal extension rescues plastid rRNA processing but not potassium homeostasis. **A)** Schematic overview of the plastid rRNA/tRNA operon and RNA intermediates and products generated during maturation. Arrowheads indicate processing sites. Binding sites of hybridization probes (A–E) used for RNA blot analyses are underlined. **B)** Detailed RNA blot analysis using probes A–E to identify plastid rRNA processing intermediates. All probes detect an increased abundance of the 3.2-kb precursor in *kea1kea2*, indicating impaired processing of the 23S–4.5S rRNA precursor. **C)** RNA blot analysis of sucrose gradient fractions using probe D against 4.5S rRNA reveals the 23S–4.5S precursor in all polysomal fractions (7–12). Higher fraction numbers correspond to heavier polysomes. Fractions 1–6 represent non-polysomal complexes, including ribosomal subunits and monosomes. **D)** RNA blot analysis using probe D demonstrates rescue of the plastid rRNA processing defect in the *KEA1ΔN* and KEA1 full-length *kea1kea2* complementation line. **E)** Potassium (K) content of leaf tissue from the indicated genotypes determined by TXRF and normalized to dry weight (DW). Boxes indicate the interquartile range (25^th^ – 75^th^ percentile), the center line represents the median, and whiskers indicate the minimum and maximum values. Statistical significance was determined by ordinary one-way ANOVA followed by Tukey’s multiple-comparison test. Different letters indicate statistically significant differences (p < 0.05; *n* = 13).

rRNA maturation is known to occur in coordination with ribosome assembly. Cleavage events generating the 4.5S rRNA and the hidden nicks within the 23S rRNA take place while the rRNA is already associated with proteins of the large (50S) ribosomal subunit. These processing steps represent some of the final stages in the formation of fully mature ribosomes (Schmid et al., 2024). We utilized the 4.5S specific probe (D) to determine whether KEA1/2 loss leads to stalled ribosomes or whether incomplete large subunits still assemble into mature ribosomes (monosomes) and subsequently into mRNA-bound ribosomes (polysomes). RNA blot analysis of polysome loading showed that the 23S – 4.5S rRNA precursor is present even in the heaviest polysome fractions, which was especially evident in *kea1kea2* plants (Fig. 4C, S6A). The same assay conducted with polysome-dissociating EDTA (instead of Mg^2+^) led to a re-distribution of the 23S – 4.5S rRNA precursor together with all other detectable rRNAs to the non-polysomal fractions, affirming its association to transcribed mRNA clusters (Fig. S6B).

To assess whether the phenotypic rescue in the complemented *KEA1ΔN* lines is accompanied by WT-like processing of the 23S – 4.5S rRNA precursor, probe D was also applied to RNA blots from plants expressing N-terminally truncated KEA1 in direct comparison to WT and *kea1kea2* complemented with full-length KEA1 (Fig. 4D). All complementation lines displayed slightly elevated levels of the unprocessed rRNA precursor compared to WT, but these levels were clearly reduced relative to the *kea1kea2* knockout (Fig. 4D). Thus, the aberrant accumulation of unprocessed rRNA observed in the mutant is largely alleviated by expression of KEA1 lacking its CC-containing N-terminal^1-479^ region.

Given that ribosome biogenesis is critically dependent on ion homeostasis (Nierhaus, 2014), which is disrupted in *kea1kea2* mutants (DeTar et al., 2021), we lastly quantified leaf element levels. For WT and *kea1kea2*, we could reproduce previous results, i.e., the double mutant displayed significantly higher potassium (K) levels than the WT (Höhner et al., 2016). Leaf K levels in the *pKEA1::KEA1-YFP* line were not significantly different from WT. In contrast, *pKEA1::KEA1ΔN* did not fully revert to WT level and exhibited significantly higher leaf K content than WT. However, K levels *pKEA1::KEA1ΔN* remained significantly lower than in *kea1kea2* (Fig. 4E). Similar changes were detected for rubidium (Rb). Notably, the differences in K and Rb levels were specific to *kea1kea2* and *pKEA1::KEA1ΔN*, whereas the majority of other elements were either not significantly altered relative to the WT or displayed only marginal variation (Table S5). These findings indicate a specific perturbation of potassium transport.

Altogether, these findings suggest that the critical K threshold necessary to disrupt rRNA processing is not reached in the *pKEA1::KEA1ΔN* line, or elevated potassium levels alone may not fully account for the accumulation of rRNA precursors.

### Conclusions

In summary, loss of IE KEAs increases the abundance of immature ribosomes, including those in polysome fractions. KEA1 is required to maintain plastid pH and K^+^ homeostasis, thereby supporting plastid gene expression (PGE) and normal growth, whereas its absence leads to elevated plastid K^+^, pH, impaired PGE, and growth defects. Notably, the KEA1 N-terminal extension has been conserved throughout the green lineage, yet KEA1ΔN largely restores PGE and growth even though plastid K^+^ accumulation persists. This unexpected result suggests that the conserved N-terminal extension modulates KEA1 transport activity *in vivo* rather than being essential for KEA1 activity per se, raising the question of why this regulatory module has been maintained over such a long evolutionary timescale. Together, these results suggest that elevated plastid K^+^ alone does not fully account for the *kea1kea2* phenotype, and that additional factors or a threshold effect likely influence plastid rRNA processing and plant growth (Fig. 5).

**Figure 5.**
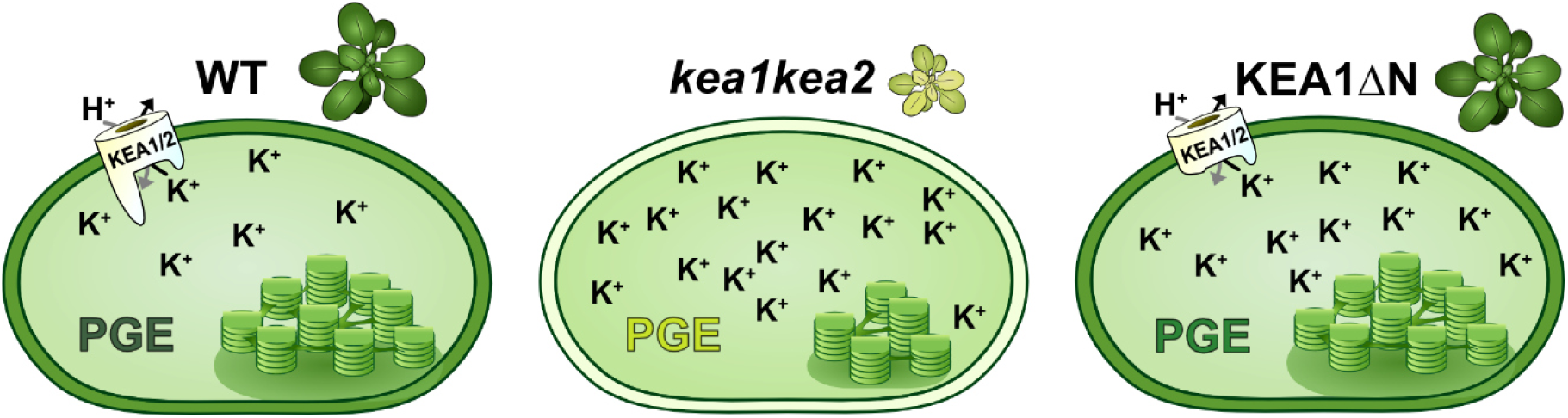
Proposed model for the role of the KEA1 N-terminal extension. WT KEA1 maintains plastid K^+^ homeostasis, enabling normal plastid gene expression (PGE) and growth. In *kea1kea2*, loss of KEA1 leads to elevated plastid K^+^ levels, impaired PGE, and reduced growth. KEA1ΔN largely restores PGE while plastid K^+^ levels remain elevated, consistent with the idea that the N-terminal extension contributes to the regulation of KEA1 transport activity *in vivo*. The observation that plastid gene expression and growth are largely restored despite elevated plastid K^+^ levels suggests that a critical K^+^ threshold may be required to impair plastid rRNA processing, or that K^+^ accumulation alone is insufficient to explain the *kea1kea2* phenotype.

## Material and methods

### Protein domain predictions and phylogenetics

Initial protein domain and structural predictions were carried out using Alphafold3 (Abramson et al., 2024), Uniprot (Consortium, 2025), IBS 2.0 (Xie et al., 2022). Subsequently, protein releases from the genomes of photosynthetic eukaryotes and cyanobacteria, shown in Table S1, were downloaded from the relevant databases and repositories and filtered for the longest isoform of each protein. Protein databases were assembled on this dataset. The six members of the KEA family from *A. thaliana* were queried against these databases via BlastP. The e-value cut-off was 1e-6. Multiple sequence alignments (MSA) were computed using MAFFT v7.505 (Katoh and Standley, 2013) with the FFT-NS-i approach and iterative refinement method. The maximum likelihood phylogeny was computed using IQ-TREE multicore version 2.0.6 (Minh et al., 2020); the best-fit model for protein evolution was determined using ModelFinder (Kalyaanamoorthy et al., 2017) as LG+G4 according to Bayesian Information Criterion (BIC), and 1,000 ultrafast bootstrapping (Hoang et al., 2018) was carried out. The clade corresponding to KEA1/2 was determined from the tree. For the proteins in this clade, proteins much shorter than the expected length of KEA1/2 homologs were removed. A new MSA was computed with the remaining proteins in this clade using MAFFT v7.505 with the FFT-NS-2 approach and progressive method. The alignment was split into two; one corresponding to the N-terminus insertion of KEA1 (61-582 aa) and the other corresponding to the rest of KEA1 (583-1193 aa). Maximum-likelihood phylogenies were computed on the split alignments using IQ-TREE as before, with ModelFinder selecting JTT+F+G4 and LG+G4 as the best-fit models for the tree of N-terminus and the tree of truncated KEA1, respectively. The topologies of these trees were compared using the phytools version 2.4-4 (Revell, 2024) in R. All the phylogenetic trees were visualized in iTOL (Letunic and Bork, 2024). The annotations were made in iTOL and Affinity Designer 2. All multiple sequence alignments were visualized in SeaView version 4 (Gouy et al., 2010).

### Plant cultivation

*A. thaliana* ecotype Col-0 (throughout all genotypes) was cultivated for all experiments on soil (Substrate A210, Stender, Schermbeck, Germany). Generally, plants were grown under long day conditions (16 h illumination / 8 h dark) at 110 µmol photons m^-2^ s^-1^, 22°C, 50% humidity.

### Stable transformation of *A. thaliana*

Full-length genomic *KEA1* (ATG to STOP codon) was inserted into derivatives of the *pGreen2.0* vector system (Pratt et al., 2020; Schwartz et al., 2025) under the control of either the constitutive pUBQ10 promoter or the endogenous *KEA1* promoter (*pKEA1*). For *pKEA1* constructs, the *pUBQ10* promoter was replaced with a 698-bp region upstream of the *KEA1* start codon. N-terminal truncation constructs were generated by Gibson assembly. Primers and cloning details are provided in Tables S2 and S3. *Arabidopsis kea1-1kea2-1* plants were transformed by the floral dip method (Clough and Bent, 1998) using the GV3101 agrobacteria strain carrying the help plasmid *pSOUP* (Hellens et al., 2000). Transgenic plants were identified on ½ MS medium containing 25 µg ml^-1^ (w/v) hygromycin. Transgene expression was verified by CLSM in the T_1_ generation and confirmed in T_2_ plants by immunoblotting using α-KEA1(2) (Bölter et al., 2020) or α-GFP antibodies (Chromotek).

### Expression and purification of N-terminal Fragments of AtKEA1 in *E. coli*

The DNA sequence encoding amino acids 135–354 of the AtKEA1 N-terminus, codon-optimized for heterologous expression in *E. coli* (Twist Bioscience, California, USA), was cloned into a Golden Gate-compatible derivative of the *pHUE* expression vector (Catanzariti et al., 2009) generating the plasmid *pHUE_AtKEA1_aa135–354*. Proteins were expressed in *E. coli* strain BL21 (DE3) by growing the cells at 37°C in Lysogeny broth medium to an OD600 of 0.8 before inducing overexpression with 0.5 mM isopropyl β-d-1-thiogalactopyranoside overnight at 20°C. Cells were lysed in 20 mM Tris-HCl pH 8.0, 300 mM NaCl, and 10 mM imidazole in the presence of 2 mM phenylmethanesulfonyl fluoride (PMSF) using a Microfluidizer® (Microfluidics™). His-tagged KEA proteins were extracted from the soluble fraction by applying a 1ml HisTRAP™ column (Cytiva). After eluting the proteins with an imidazole gradient, the N-terminal His6–ubiquitin moiety was cleaved (Baker 2005). Finally, proteins were separated by size-exclusion chromatography (HiLoad® 16/60 Superdex® 200 PG, GE Healthcare) in 20 mM Tris-HCl pH 8.0 and 50 mM NaCl. Fractions of interest were concentrated for crystallization screening.

### Immunoblotting

Immunoblotting was carried out either on total leaf tissue (Völkner et al., 2021; Völkner et al., 2024) or on total protein of isolated chloroplasts. For chloroplasts, plastids were collected and resuspended in 1x SDS-loading dye (1% (v/v) glycerol, 5% (v/v) β-mercaptoethanol, 2% (w/v) SDS, 62.5 mM Tris-HCl pH 6.8, 0.003% (w/v) bromophenol blue) and heated at 95°C for 5 min. In case of leaf tissue, equal amounts of leaf tissue were homogenized and subsequently solubilized in 10% (v/v) glycerol, 150 mM NaCl, 2 mM EDTA, 5 mM DTT, 50 mM Tris-HCl pH 7.5, 1% (w/v) SDS, plant protease inhibitor mix for 30 min at 4°C. Subsequently the isolates were centrifuged at 16.000*g* for 10 min and the supernatant was mixed with SDS-loading dye and boiled at 95°C for 5 min. Proteins were separated on a 10.5% (w/v) - 15% (w/v) SDS-PAGE depending on the molecular weight of the protein of interest and transferred to a PVDF membrane using a wet-blotting system (transfer buffer: 25 mM Tris/HCl, pH 8.3, 192 mM glycine, 20 % (v/v) MeOH, 0.1 % (w/v) SDS). Immunodetection was carried out with the respective antibody at 4°C overnight in TBS-T buffer (50 mM Tris/HCl pH 7.5, 150 mM NaCl, 0.05 % Tween20) (KEA1: (Bölter et al., 2020), KEA3: (Armbruster et al., 2014), GFP or RFP (ChromoTek, Polyclonal antibody)). Target proteins were visualized using enhanced chemiluminescence by anti-rabbit secondary antibody fused to horse radish peroxidase (Merck KGaA, Darmstadt, Germany).

### Localization of KEA1 in *A. thaliana* protoplasts

Initially, protoplasts from transgenic reporter lines were prepared using the rapid sandwich prep method (Wu et al., 2009). Detailed localization studies were carried out on a Leica Stellaris 5 (Wetzlar, Germany) as described before (Höhner et al., 2019). In brief, YFP and chlorophyll colocalization was done using 514 nm for excitation. Emission was collected with hybrid detectors at 518–565 nm and 627–706 nm for YFP and chlorophyll autofluorescence, respectively.

### Blue Native (BN) PAGE

BN-PAGE on isolated chloroplasts was performed as reported by (Völkner et al., 2021). In brief, isolated chloroplasts equal to 15 µg chlorophyll per sample were solubilized for 15 min on ice using 0.5% (w/v) dodecyl maltoside. Subsequently, samples were mixed with BN-loading dye (750 mM aminocaproic acid, 5% (w/v) Coomassie G-250) and separated on a 5 – 15% (v/v) native acrylamide gradient. For detection of the target protein, the BN PAGE was incubated in transfer buffer with increased SDS content (25 mM Tris/HCl, pH 8.3, 192 mM glycine, 20 % (v/v) MeOH, 1 % (w/v) SDS). Subsequently, transfer of proteins and detection of the target protein was carried out as described for immunoblotting.

### Size exclusion chromatography (SEC) analysis of solubilized chloroplasts

Intact chloroplasts equivalent to 250 µg chlorophyll were solubilized using SEC buffer (10 mM phosphate buffer pH 7.0, 2.7 mM KCl, 137 mM NaCl, 10 mM EDTA/NaOH) containing 0.5 % Triton X-100. Solubilization was carried out as described for the BN-PAGE samples. Solubilized chloroplasts were separated on a Superose 6 column (Cytiva) using SEC buffer containing 0.05% Triton X-100. Elution fractions were immunoblotted using a KEA1(2) or KEA3 specific immunoglobulin, respectively. To detect the well described RuBisCO complex, elution fractions were separated on an SDS-PAGE and stained with Coomassie brilliant blue (CBB). For size comparison, commercial high molecular weight standards (blue dextran, thyroglobulin, ferritin, aldolase; GE Healthcare) were separated on the same column using SEC buffer which did not contain Triton X-100. Elution of the respective standards was detected by UV absorption at 280 nm.

### STED Microscopy

Plants were grown for 18 days in a long day growth regime. The younger leaves were used for protoplast isolation and processed as described (Legen et al., 2024). After fixation and embedding, microscopy was done on an Abberrior Facility Line fluorescence microscope using a 515 nm laser for YFP excitation and a 405 nm laser for the DAPI excitation. The latter was also used to excite chlorophyll autofluorescence. The same settings for all three channels were used on the KEA1-YFP and WT protoplasts. Images for all four channels (Autofluorescence, DAPI, Venus and bright field) were taken simultaneously. Distance measurements between nucleoids (DAPI channel) and KEA1 puncta (YFP channel) were done as described previously (Legen et al., 2024).

### GFP affinity purification

Isolated chloroplasts or nitrogen ground plant leaf material was solubilized in protein isolation buffer (10% (v/v) glycerol, 150 mM NaCl, 2 mM EDTA, 50 mM Tris-HCl pH 7.5, 0.5 % (w/v) Triton X-100, plant protease inhibitor mix) for 15 min on ice followed by centrifugation of 16.000*g* for 10 min. For immunoprecipitation of mVenus-tagged proteins a homemade GFP-trap was added to the supernatant and incubated in an overhead shaker for 2 – 4 h (Schwartz et al., 2025). Subsequently, the GFP trap was washed three times to remove unbound proteins. For analysis of co-immunoprecipitated proteins the GFP-trap was heated with 1x SDS loading dye (1% (v/v) glycerol, 5% (v/v) β-mercaptoethanol, 2% (w/v) SDS, 62.5 mM Tris-HCl pH 6.8, 0.03‰ (w/v) bromophenol blue) and heated at 95°C for 5 min. Subsequently, proteins were separated on an SDS-PAGE and detected via immunoblotting.

### Transient transformation of *N. benthamiana*

Transient double transformation of *N. benthamiana* was carried out as described by (Waadt and Kudla, 2008). In brief, Agrobacteria were cultivated to OD_600_ of ∼3 in LB containing the respective antibiotics for selection. Subsequently, agrobacteria were collected and resuspended to an OD600 of 0.4 in infiltration media (10 mM MES, pH 6.0, 10 mM MgCl_2_, 150 µM acetosyringone). For double infiltration, two constructs were combined. For each single and double injection, agrobacteria containing the p19 were added to OD_600_ of 0.2. After incubation for 4 h in the dark *N. benthamiana* plants were infiltrated with the respective constructs using a syringe.

### Acquisition of photosynthetic parameters

Plants were dark-adapted for 20 min. Subsequently, chlorophyll *a* fluorescence was measured using standard light curve setting on a Walz IMAGING-PAM M-Series MAXI version (Walz, Effeltrich, Germany). False-color images were exported using the ImagingWinGigE software. Data were analyzed as described previously (Schneider et al., 2019).

### *E. coli* complementation

Codon-optimized cDNA coding for full length KEA1 without the transit peptide was synthesized by TWIST Bioscience (San Francisco, CA, USA) and served as the PCR template to generate the truncated construct. Gene fragments were inserted via NcoI/XhoI restriction sites followed by Ligation with T4 Ligase into a modified pACBAD14 vector cut with the same enzymes in which the original chloramphenicol resistance gene had been replaced with AmpR, a beta-lactamase, that confers ampicillin and carbenicillin resistance in bacteria. In addition, the vector was outfitted with a C-terminal mVENUS for protein fusion.

*E. coli* LB2003 (F-, *thi, lacz, gal, rha, ΔkdpFABC5, trkD1, ΔtrkA*) was transformed with pBAD-KEA1-YFP, pBAD-KEA1ΔN^480–1193^-YFP and pBAD-YFP plasmids and grown overnight in high K^+^ medium (LBK) containing 5 g l^-1^ yeast extract, 10 g l^-1^ tryptone, 150 mM KCl in the presence of 100 mg l^-1^ carbenicillin. From the overnight culture, 100 μl was diluted in 5 ml of LBK and grown to OD_600_ = 0.5. Expression was induced by the addition of 0,02‰ arabinose. After 2 h of induction, the cell density was measured, and the culture was diluted (normalized) to OD_600_ 0.5. Four-μl drops of the diluted cell suspension were spotted on carbenicillin plates containing 5 g l^-1^ yeast extract, 10 g l^-1^ tryptone, and either 0 or 150 mM KCl with arabinose and carbenicillin. Reproducibility of complementation assays was confirmed in several independent replicate experiments (n=3).

### Polysome loading analysis and RNA gel blot hybridization

Polysome loading experiments were conducted as described recently (Manavski et al., 2025) following the protocol outlined by (Barkan, 1993). To induce polysome release/dissociation, puromycin and EDTA were used, as described in (Barkan, 1993). Total RNA extraction, followed by gel separation, blotting, hybridization, and signal detection were performed as previously detailed (Manavski et al., 2021; Wunder et al., 2026). 80-mer DNA oligos probes (Table S2) were 5′ end-radio-labeled using T4 polynucleotide kinase (NEB, Ipswich, MA, USA) according to the manufacturer’s protocol or 3’-Digoxigenin (DIG)-labeled DNA probes from Metabion international AG, Germany.

### Quantification of leaf elements

Leaf tissue elemental composition was determined by total X-ray fluorescence (TXRF) as initially described by (Holzner et al., 2026). Initially, leaf tissue samples were dried at 60°C and homogenized using a zirconium mortar (Stanford Advanced Materials, Santa Ana, CA, USA). Aliquots of 3 mg dry powder were digested in 200 µl of 69% (v/v) HNO₃ at 95°C for 90 min. Following digestion, samples were combined with element standards at a 1:10 ratio to final concentrations of 1 ppm vanadium (WL) and 1 ppm gallium (Mo-K). A 69% (v/v) HNO₃ blank was measured to determine background levels.

All samples were applied onto siliconized quartz glass carriers (B&M Optik, Pirna, Germany) and analyzed at 50 kV using Mo-K and WL excitation, with 600 s acquisition time on a S4 T-STAR instrument (Bruker, Berlin, Germany) . Elemental concentrations were normalized dry weight. Background contributions, determined from the respective buffer or acid blanks, were subtracted from all measurements. If backgorund levels were higher than the measured element levels, the respective element was considered as non-detectable (ND). Statistical analyses were performed using an ordinary one-way analysis of variance (ANOVA) followed by Tukey’s multiple comparison test (GraphPad Prism 10.4.1, Dotmatics, Boston, USA). Differences were considered statistically significant at P < 0.05.

### Crystallization

The crystallization experiments for *At*KEA1 135-354 fragment were performed at the X-ray Crystallography Platform at Helmholtz Munich, Germany. The best diffracting crystal was obtained from 0.056 M NaH_2_PO^4^, 1.344 M K_2_HPO_4_ (pH 8.2). For the X-ray diffraction experiments, crystal was mounted in the nylon fiber loop and flash-cooled to 100 K in liquid nitrogen. The cryoprotection was performed for a few seconds in reservoir solution complemented with 25% (v/v) ethylene glycol. Diffraction data was collected on the PETRAIII P11 beamline. The data was indexed and integrated using *XDS* (Kabsch, 2010) and scaled using *SCALA* (Evans, 2006; Winn et al., 2011). Intensities were converted to structure-factor amplitudes using the program *TRUNCATE* (French and Wilson, 1978). Supplementary Table S4 summarizes data collection and processing statistics.

### Structure determination and refinement

The structure of *the AtKEA1* 135-354 fragment was solved by molecular replacement using the model generated by *SwissModel* (Waterhouse et al., 2018). For molecular replacement calculations, the program *MolRep (Vagin and Teplyakov, 1997; Winn et al., 2011)* was used. Model rebuilding was performed with *COOT* (Emsley and Cowtan, 2004). The refinement was done with *REFMAC5* (Murshudov et al., 1997) using the maximum-likelihood target function. The current model is characterized by R and R_free_ factors of 37.5/40%, respectively. The stereochemical analysis of the final model was done with *PROCHECK* (Laskowski et al., 1993). For all the crystallographic calculations, the *SBGrid* software bundle was used (Morin et al., 2013). The refinement statistics for the current model are summarized in Supplementary Table S4. Due to low quality of the data the structure has not been deposited to the Protein Data Bank.

### Accession numbers

*KEA1* (At1g01790), *KEA2* (At4g00630), *KEA3* (At4g04850), *kea1-1kea2-1* (SAIL_586_D02, SALK_045324, ABRC: CS72318 or NASC: N72318) (Kunz et al. 2014), *kea3-1* (SAIL_556_E12, ABRC: CS72321 or NASC: N72321). pG20_mVenus_Hyg (GenBank: MT896404.1, Addgene Plasmid #159703)

## Acknowledgments

We thank Dr. Nobuyuki Uozumi (Tohoku University) for providing the LB2003 strain. We acknowledge Dr. Martin K. M. Engqvist (Epoch Biodesign) for providing pACBAD14 and Dr. Geoffry Davis (LMU Munich) for modifying it. We thank Benjamin Kouri, Beata Szulc, and gardener Albert Schorer (all at LMU Munich) for excellent technical support. Lastly, we thank Alexander Schober from NTU Singapore for providing the Goldengate compatible pHue vector.

## Author contributions

T.W., B.B., and H.-H.K. conceived the study and wrote the manuscript. T.W. generated constructs, isolated plant mutants, performed experiments, analyzed data, and prepared figures. L.J.H. conducted BN and TXRF analyses, evaluated elemental data, and contributed to manuscript writing. B.B. generated constructs for bacterial complementation, isolated plant mutants, performed experiments, analyzed data, and prepared figures. N.M. and J.M. assisted with polysome experiments, while N.M. and J.F. performed RNA blot analyses. C.F.K., M.N.B., and J.dV. conducted homology investigations, phylogenetic analyses, evolutionary inferences and contributed to the corresponding manuscript section. J.L. performed STED microscopy, with data analysis by F.R. S.M. carried out confocal imaging, and R.J. and D.N. performed crystallization and data analysis. F.H. was involved in data analysis. All authors contributed to manuscript revision.

