## Supplemental Figures for "The Conserved N-Terminal Extension of AtKEA1 Is Largely Dispensable for Plastid Function but Contributes to Potassium Homeostasis"

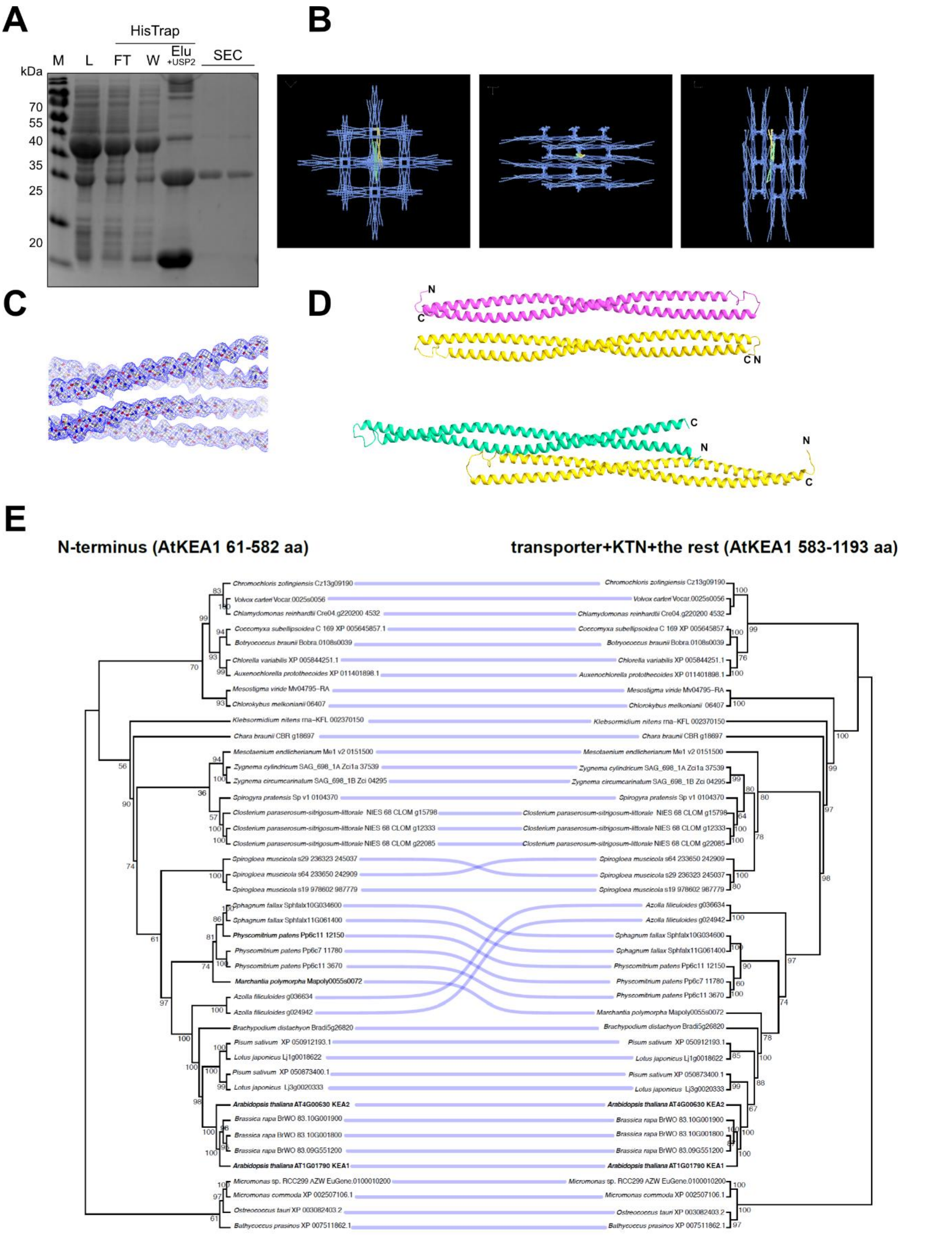


**Figure S1. Crystallization of AtKEA1's N-terminal coiled-coil domain and phylogeny of the two KEA1 components. A)** Coomassie-stained SDS–PAGE gel showing purification of the N-terminal AtKEA1 coiled-coil fragment (aa 135–354) before crystallization. M, molecular weight marker; L, lysate; FT, HisTrap flow-through; W, HisTrap wash; Elu+USP2, USP2 protease-treated HisTrap elution; GF, Gel filtration fractions. **B)** Crystal packing of the AtKEA1 aa 135–354 fragment viewed along crystallographic axis *c* (*left*), axis *b* (*middle*), and axis *a* (*right*). The two molecules in the asymmetric unit are shown in green and yellow, whereas symmetry-related molecules are shown in blue. **C)** 2Fo-Fc electron density map contoured at 1σ for a representative region of AtKEA1's coiled-coil structure. **D)** Packing reveals two potential dimeric arrangements: antiparallel (*top*) and parallel (*bottom*). Individual coiled-coil monomers are shown in different colors. **E)** Co-phylogeny plot for pairwise comparison of the trees for the N-terminal region containing the crystallized coiled-coil domain (left) versus the residual membrane-bound KEA1 fragment that harbors the transporter and the KTN domains. Plot was generated using ape and phytools packages in R. Trees were rooted at the midpoint.

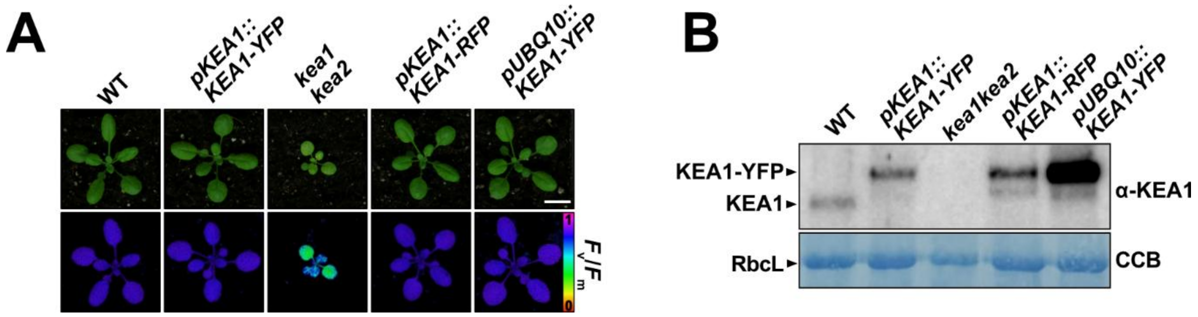


**Figure S2. Fluorescently tagged full-length KEA1 driven by either the native or the UBIQUITIN10 (UBQ10) promoter complements the *kea1kea2* phenotype. A)** Representative RGB images and corresponding false-color $F_v/F_m$ images of WT, the *kea1kea2* null mutant, and complementation lines expressing KEA1-YFP under the control of the native KEA1 promoter (*pKEA1::KEA1-YFP*) or the constitutive UBQ10 promoter (*pUBQ10::KEA1-YFP*) in the *kea1kea2* background. Scale bar = 1 cm. **B)** Immunoblot using the α-KEA1 antibody to detect KEA1 expression in the complementation lines. A Coomassie-stained membrane (CCB) showing the Rubisco large subunit (RbcL) is included as a loading control.

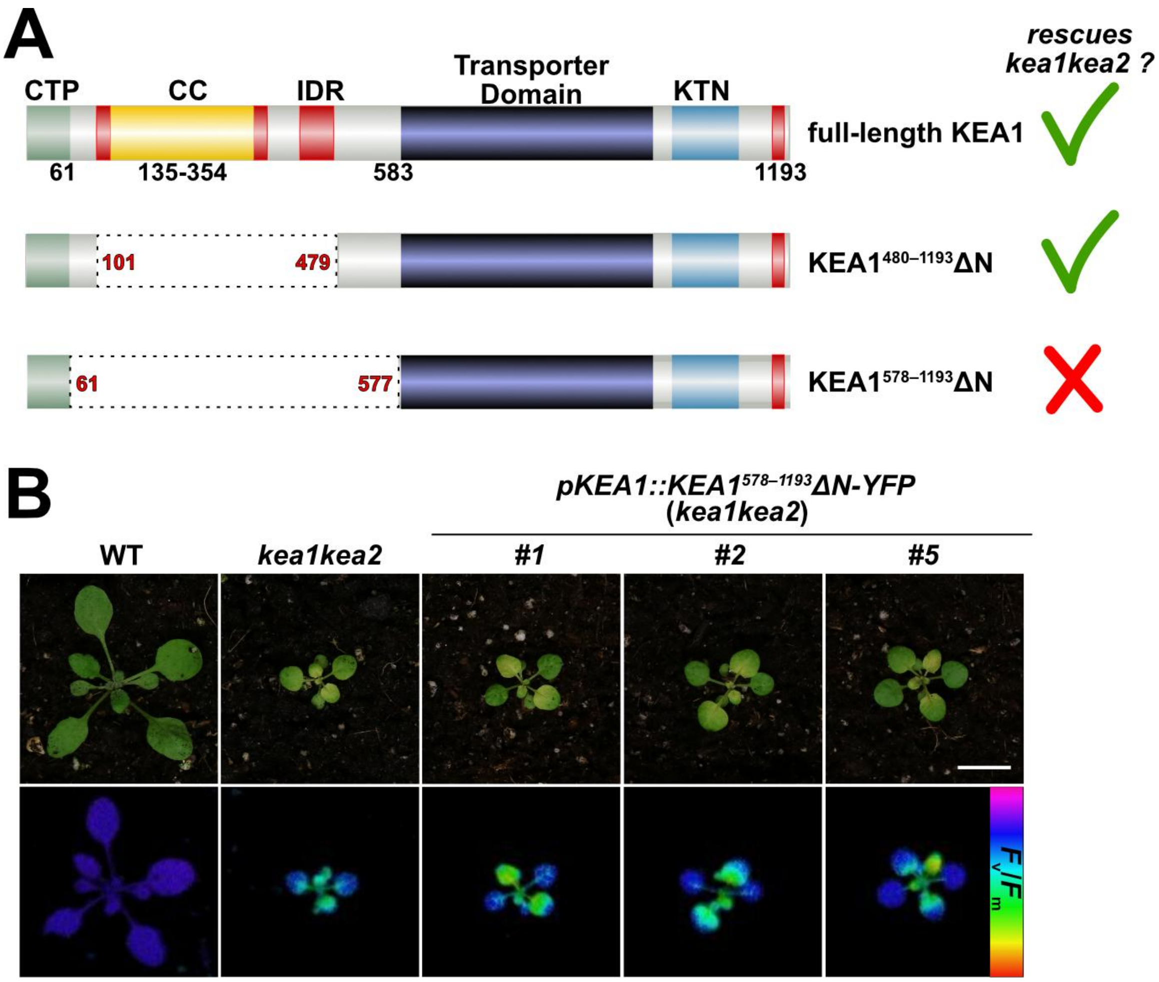


**Figure S3. A further N-terminal truncation of KEA1 fails to complement the *kea1kea2* phenotype. A)** Schematic representation of full-length KEA1 and the two N-terminal truncation variants used for complementation of *kea1kea2*. The functional KEA1$^{480–1193}$ΔN variant retains amino acids 1–100 fused directly to amino acids 480–1193. The further truncated KEA1$^{578–1193}$ΔN variant retains only amino acids 1–60 fused directly to amino acids 578–1193. Dashed lines indicate deleted regions. CTP, chloroplast targeting peptide; CC, coiled-coil domain; IDR, intrinsic disordered regions; KTN, Potassium Transport–Nucleotide binding domain. **B)** Representative RGB images and corresponding false-color $F_v/F_m$ images of WT, the *kea1kea2* knockout, and transgenic lines expressing KEA1$^{578–1193}$ΔN-YFP under the control of the native KEA1 promoter (*pKEA1::KEA1$^{578–1193}$ΔN-YFP*). Transgenic plants were selected on ½ MS medium containing 25 µg ml$^{-1}$ (w/v) hygromycin and transferred to soil after 2 weeks. Scale bar = 1 cm.

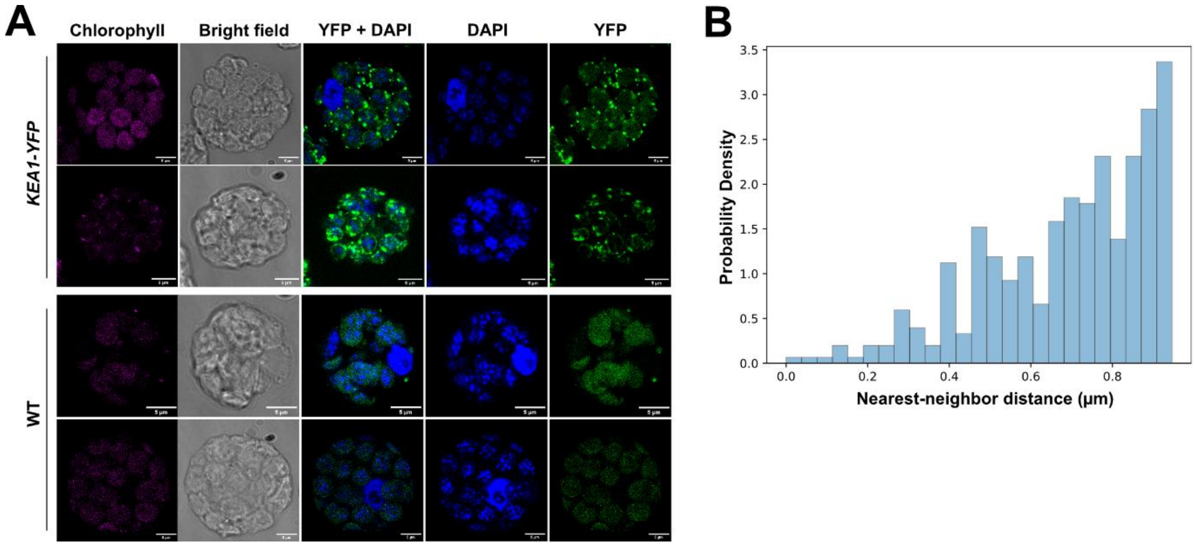


**Figure S4. KEA1 does not associate with plastid nucleoids. A)** STED microscopy of DAPI-stained protoplasts isolated from WT and complemented KEA1-YFP plants. Chlorophyll autofluorescence, bright-field, merged YFP/DAPI, DAPI, and YFP channels are shown. Scale bar = 5 µm. **B)** Distribution of nearest-neighbor distances between DAPI-labelled plastid nucleoids and KEA1-YFP puncta. For each DAPI signal maximum, the distance to the nearest KEA1-YFP signal maximum was determined. The normalized histogram is based on approximately 400 distance measurements obtained from up to 23 z-slices of 12 independent samples.

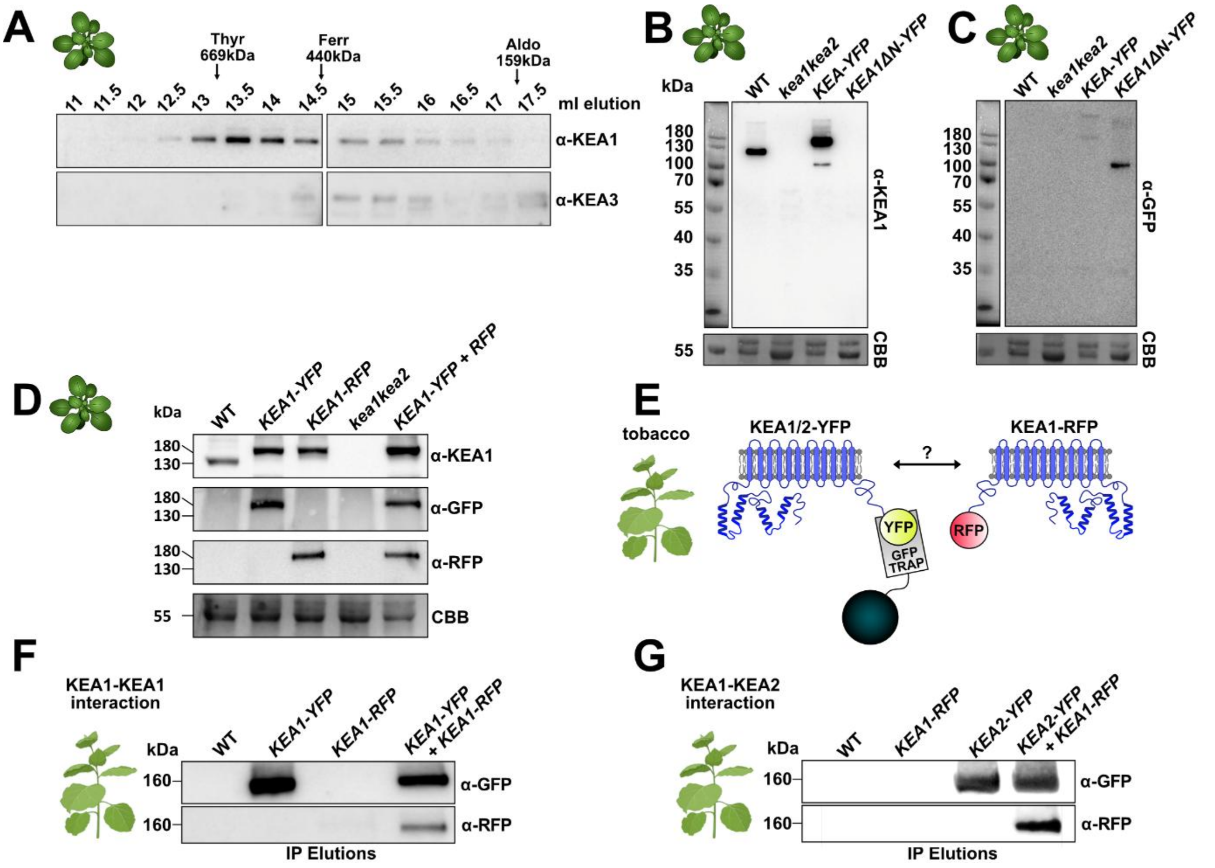


**Figure S5. KEA1 self-association and interaction with KEA2. A)** Size-exclusion chromatography of detergent-solubilized WT chloroplasts followed by immunoblot analysis of KEA1 and KEA3 in the indicated elution fractions. Elution volumes of molecular weight standards are indicated above the fractions. **B, C)** Immunoblot analysis of samples corresponding to the Blue Native-PAGE experiment shown in Fig. 3B following SDS-PAGE using antibodies against KEA1 (B) and GFP (C). Coomassie Brilliant Blue (CBB) staining is shown as a loading control. **D)** Immunoblot analysis of total protein extracts from the indicated stable *Arabidopsis* lines using antibodies against KEA1, GFP, and RFP. CBB staining is shown as a loading control. These lines were used for the GFP affinity purification experiment shown in Fig. 3E. **E)** Schematic of the transient expression assay performed in *Nicotiana benthamiana*. KEA1-YFP or KEA2-YFP was co-expressed with KEA1-RFP under the control of the constitutive UBQ10 promoter, and the YFP-tagged protein was purified using GFP-Trap beads (YFP is recognized by GFP-Trap). **F)** Immunoblot analysis of GFP-Trap eluates demonstrating co-purification of KEA1-RFP with KEA1-YFP, consistent with KEA1 self-association. **G)** Immunoblot analysis of GFP-Trap eluates demonstrating co-purification of KEA1-RFP with KEA2-YFP, consistent with an interaction between KEA1 and KEA2.

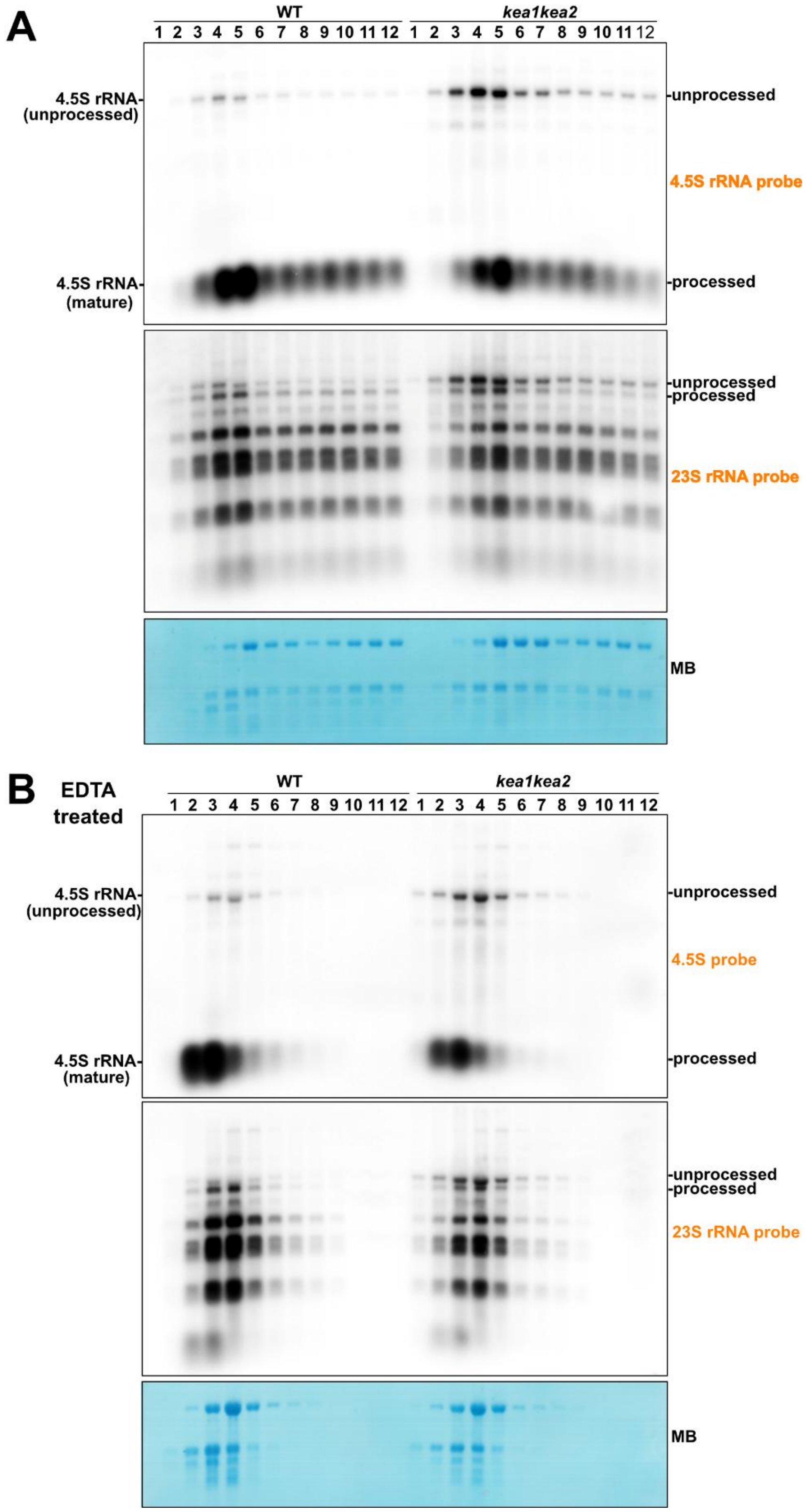


**Figure S6. rRNA precursors are incorporated into plastid ribosomes and polysomes.** Polysomes were isolated from WT and *kea1kea2* plants in the presence of $Mg^{2+}$ (**A**) to preserve polysomes or EDTA (**B**) to dissociate ribosomes. Sucrose gradient fractions were analyzed by RNA blot using probes specific for the 4.5S and 23S rRNAs. Higher fraction numbers correspond to heavier polysomes. Methylene blue (M.B.) staining was used to visualize total RNA. Panel A is reproduced in part in Fig. 4C.
